# Different Munc18 Proteins Mediate Baseline and Stimulated Airway Mucin Secretion

**DOI:** 10.1101/451914

**Authors:** Ana M. Jaramillo, Lucia Piccotti, Walter V. Velasco, Anna Sofia Huerta Delgado, Zoulikha Azzegagh, Felicity Chung, Usman Nazeer, Junaid Farooq, Josh Brenner, Jan Parker-Thornburg, Brenton L. Scott, Christopher M. Evans, Roberto Adachi, Alan R. Burns, Silvia M. Kreda, Michael J. Tuvim, Burton F. Dickey

**Affiliations:** Department of Pulmonary Medicine, The University of Texas MD Anderson Cancer Center, Houston, Texas, 77030, USA.; Institute of Bioscience & Technology, Texas A&M University Health Science Center, Houston, Texas, 77030, USA.; Tecnologico de Monterrey, Escuela de Medicina y Ciencias de la Salud, Monterrey, N.L., Mexico, 64710, Mexico.; Marsico Lung Institute/Cystic Fibrosis Center, University of North Carolina at Chapel Hill, Chapel Hill, North Carolina, 27599, USA.; Department of Genetics, The University of Texas MD Anderson Cancer Center, Houston, Texas, 77030 USA.; Division of Pulmonary Sciences and Critical Care Medicine, University of Colorado Denver School of Medicine, Aurora, Colorado, 80045, USA.; College of Optometry, University of Houston, Houston, Texas 77204, USA.

## Abstract

Airway mucin secretion is necessary for ciliary clearance of inhaled particles and pathogens, but can be detrimental in pathologies such as asthma and cystic fibrosis. Exocytosis in mammals requires a Munc18 scaffolding protein, and airway secretory cells express all three Munc18 isoforms. Using conditional airway epithelial deletant mice, we found that Munc18a has the major role in baseline mucin secretion, Munc18b has the major role in stimulated mucin secretion, and Munc18c does not function in mucin secretion. In an allergic asthma model, Munc18b deletion reduced airway mucus occlusion and airflow resistance. In a cystic fibrosis model, Munc18b deletion reduced airway mucus occlusion and emphysema. Munc18b deficiency in the airway epithelium did not result in any abnormalities of lung structure, particle clearance, inflammation, or bacterial infection. Our results show that regulated secretion in a polarized epithelial cell may involve more than one exocytic machine at the apical plasma membrane, and that the protective roles of mucin secretion can be preserved while therapeutically targeting its pathologic roles.

## Introduction

In mammalian conducting airways, mucus forms a critical barrier that protects the lungs from inhaled particles, pathogens and toxicants (1). These foreign substances are trapped by mucus, which is swept out of the lungs by ciliary beating into the pharynx where it is swallowed. Secreted polymeric mucins, the principal macromolecular components of mucus, are large, highly glycosylated proteins that polymerize into linear chains and networks (2, 3). Mucins are packaged dehydrated in secretory granules, and after exocytosis they interact with several hundred-fold their mass of water to expand and generate viscoelastic, gel-like mucus.

Two polymeric secreted mucins are expressed in the airway epithelium—Muc5b and Muc5ac. Mouse Muc5b is expressed constitutively in superficial epithelial cells and submucosal glands, and is primarily responsible for mucociliary clearance. Deletion of the gene encoding Muc5b in mice results in death from bacterial infection and airway obstruction (4). Heterozygous gene deletion results in ~50% reduction in polystyrene bead clearance (5), showing that Muc5b is limiting for mucociliary clearance. Conversely, an overexpressing allele of human *MUC5B* is highly prevalent in Caucasians and shows evidence of positive selection, probably for its value in protection against lung infection even though it is a risk factor for pulmonary fibrosis late in life (6, 7). Muc5ac is expressed only at low levels in all airways of naïve (uninflamed) mice and in distal airways of humans (1). However, *Muc5ac* expression rises ~40-fold during allergic inflammation (8, 9). Induced *Muc5ac* expression contributes importantly to helminth defense in the gut (10), and may help trap helminths migrating through the lungs (11). In allergic asthma, overexpressed and rapidly secreted Muc5ac causes airway mucus occlusion and airflow obstruction (12).

Mucins are secreted at a low baseline rate and a high agonist-stimulated rate (13, 14). Both rates are regulated by the second messengers diacylglycerol and calcium acting on the exocytic sensor Munc13-2 (15). Important extracellular agonists promoting baseline secretion are ATP and its metabolite adenosine, released predominantly from ciliated cells sensing shear stress from airflow during ventilation (14, 16, 17). These agonists act on heptahelical receptors coupled by G-proteins of the Gq subtype to PLC-β that generates the second messengers diacylglycerol and inositol triphosphate, with the latter inducing the release of calcium from intracellular stores (13). Higher levels of the same agonists can stimulate high rates of mucin secretion (18), as can the neural and inflammatory mediators acetylcholine and histamine acting on the same pathway downstream of their cognate receptors (12). At high levels of intracellular calcium, the fast, low-affinity exocytic calcium sensor Synaptotagmin-2 promotes mucin secretion (19). Baseline secretion is thought to be primarily responsible for clearance of inhaled particles and pathogens, while stimulated secretion can induce airway obstruction protectively to trap helminths or pathologically in asthma (11, 12).

Defects in mucin secretion in SNAP23 and VAMP8 mutant mice implicate the highly conserved SNARE (soluble N-ethylmaleimide-sensitive factor attachment protein receptors) machinery in mucin exocytosis (20, 21). The SNARE complex is a four-helix bundle comprised of three helices attached to the target membrane (t-SNAREs) and one attached to the vesicle membrane (v-SNARE). Specific binding of these helices confers accuracy and directionality on the fusion reaction, and full coiling provides the energy to fuse the lipid membranes. SNARE-dependent vesicle traffic universally involves SM (Sec1/Munc18) proteins that promote SNARE complex assembly and help prevent off-target interactions (22). Yeast contain four SM proteins, among which the exocytic protein Sec1 has evolved into three exocytic Munc18 isoforms in metazoans. We previously found, using heterozygous hypomorphic mutant mice (23), that Munc18b has a role in stimulated mucin secretion, but were unable to identify a role in baseline secretion. Here, by performing a comprehensive analysis of airway epithelial deletants of all three Munc18 isoforms in mice, we sought to identify the Munc18 protein(s) mediating baseline secretion, to fully characterize the role of Munc18b in stimulated mucin secretion, and to test the hypothesis that selective impairment of stimulated secretion can protect against airway mucus obstruction in pathophysiologic models.

## Results

### Generation of airway epithelial Munc18 deletant mice

We had previously found that homozygous mutant Munc18b mice are not viable postnatally (23), and others found that homozygous Munc18a (24) and Munc18c (25) knockout mice are similarly not viable. Therefore we assembled a panel of conditional mutant Munc18 mice by generating a conditional allele of Munc18b that is described here (Figure 1, A and S1), generating a conditional allele of Munc18c that is described elsewhere (26), and obtaining from others a conditional allele of Munc18a (27). Crossing conditional Munc18b mice with Zp3-Cre transgenic mice to generate whole animal knockout mice did not yield any Munc18b^-/-^ pups, confirming that Munc18b is an essential gene in mice. Munc18b^-/-^ embryos grown in vitro developed to E3.5 at a Mendelian ratio, but in vivo, Munc18b^-/-^ embryos at E10.5 were present at less than a Mendelian ratio, and at E11.5 there were none (Figure 1, B).

**Figure 1.**
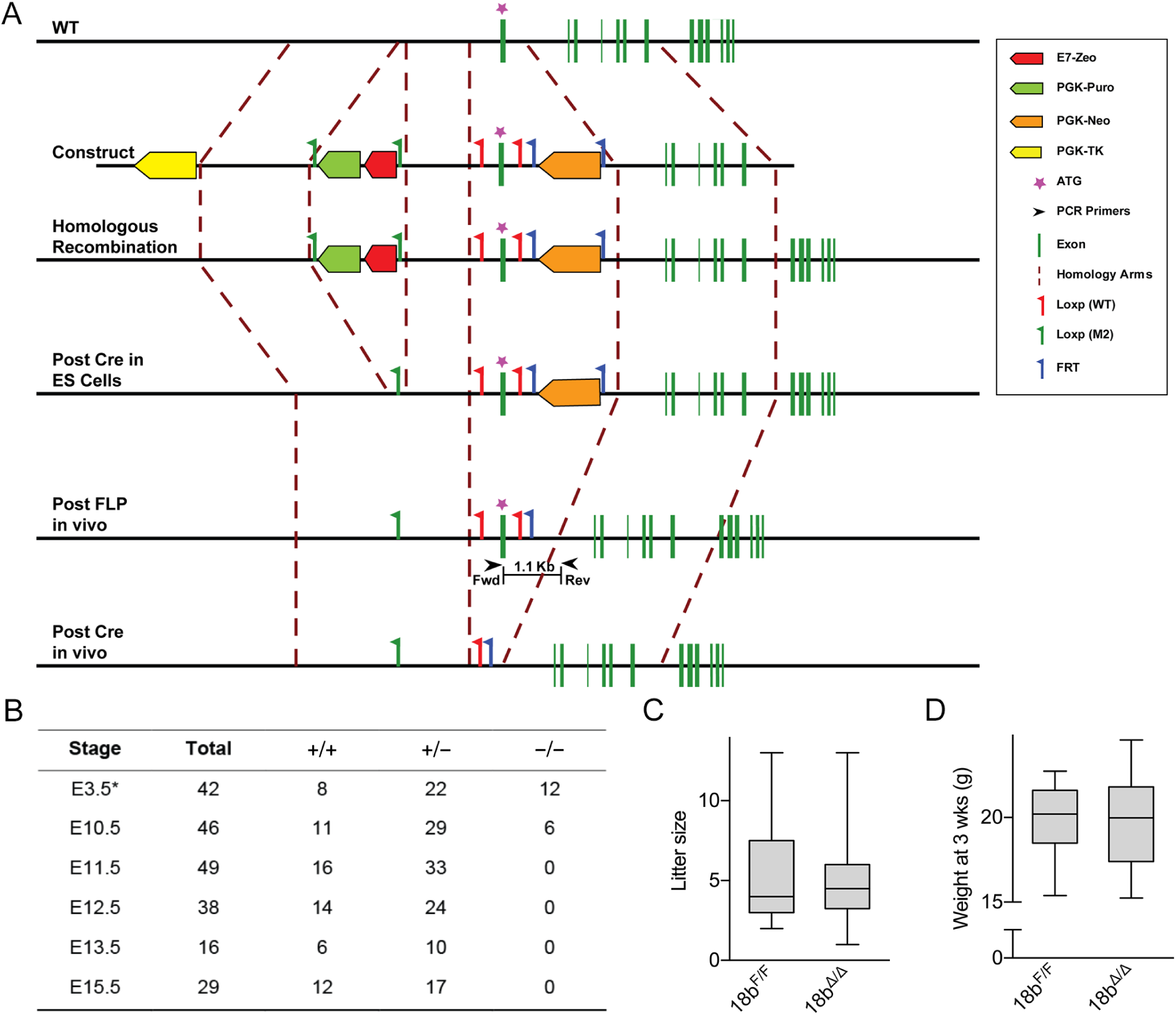
Generation of Munc18b conditional deletant mice. **(A)** Exon 1 of the Munc18b gene was flanked by two loxP sites (red flags) via homologous recombination. Herpes simplex virus thymidine kinase (yellow) was used for negative selection. Zeocin (red) and puromycin (green) resistance genes flanked by two loxP (M2) sites (green flags), removed by Cre in embryonic stem cells, and a neomycin resistance gene (orange) flanked by two FRT sites (blue flags), removed by Flp recombination, were used for positive selection. Exon 1 was removed by Cre recombination in mice. **(B)** Table showing embryos of different genotypes harvested at various days of embryonic (E) development after generating a full KO by crossing the mouse in (A) to a Zp3-Cre mouse. **(C)** Litter size after crossing floxed mouse in (A) to a CCSP^iCre^ mouse to generate a conditional deletant mouse specific for airway epithelium are compared to their floxed littermates (n=50-52 per group). **(D)** Weight at 3 weeks of age by Munc18b conditional deletants and their floxed littermates (n=50-52 per group).

Deletion of Munc18b in airway epithelial cells by crossing Munc18b^F/F^ mice with CCSP^iCre^ mice (28) yielded Munc18b^CCSP-Δ/Δ^ (hereafter Munc18b^Δ/Δ^) mice with normal litter sizes and weight at 3 weeks compared to Munc18b^F/F^ mice (Figure 1, C and D). The efficiency of recombination of CCSP^iCre^ mice at the ROSA26 locus is >99% in both airway secretory and ciliated cells (Figure S2, A and B), and occurs occasionally in alveolar type 2 secretory epithelial cells as well (Figure S2, C). Histopathologically, the lungs of all floxed and single Munc18 isoform airway deletant mice were unremarkable by H&E staining, as were the lungs of Munc18a/b and Munc18b/c floxed mice, and Munc18a/b double deletant mice (Figure S3, A and B). However, the airways of Munc18b/c double deletant mice showed a flattened epithelium with an almost complete absence of airway secretory cells by immunohistochemical staining for club cell secretory protein (CCSP) (Figure S3, C), and the lung alveolar regions showed emphysema (Figure S3, D). To determine whether these abnormalities reflected dependency on Munc18b and Munc18c only during development, we used CCSP^CreER^ mice to induce recombination during adulthood (29). Airway secretory cell viability was still impaired because CCSP expression was lost two weeks after recombination (Figure S3, E), though emphysema was not present (Figure S3, F).

To determine the normal expression of Munc18 isoforms in the airway epithelium and confirm the efficiency of gene deletion, we performed quantitative *in situ* hybridization with riboprobes. Munc18a and Munc18b transcripts were expressed in secretory cells at levels several-fold higher than in ciliated cells (Figure 2, A and B), whereas Munc18c transcripts were expressed in both cell types at similar levels (Figure 2, C). All three Munc18 conditional deletant mice showed no significant expression of cognate transcripts (Figure 2), and there was no significant difference in transcript expression between any of the floxed mice and WT (not shown).

**Figure 2.**
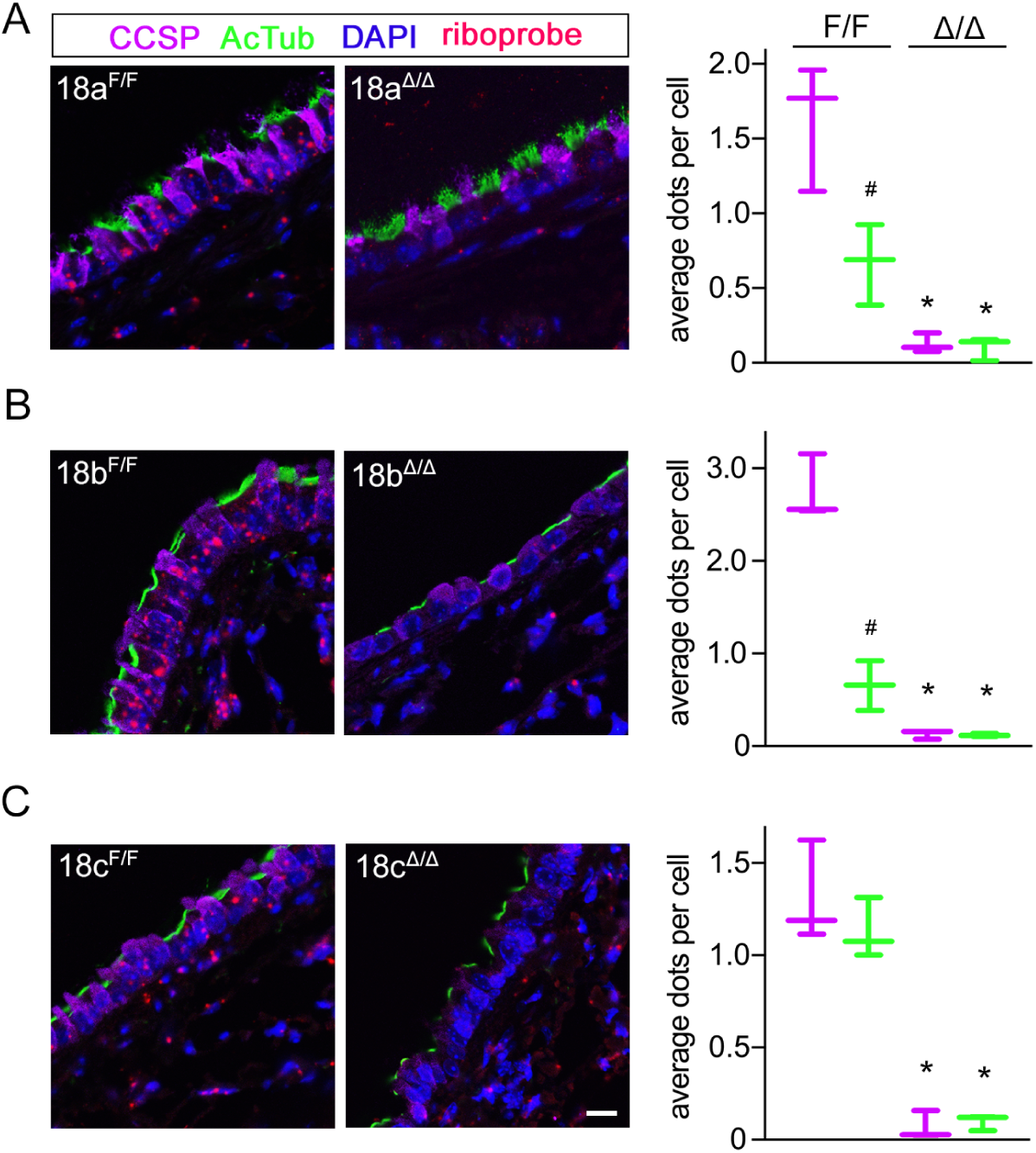
In situ hybridization of Munc18 isoforms. Representative images of in situ hybridization with fluorescent-labeled riboprobes (shown in red), specific for murine Munc18a **(A)**, Munc18b **(B)** and Munc18c **(C)**. Sections are from naïve (uninflamed) lungs of floxed and conditional deletant mice. CCSP was used as a secretory cell marker (purple) and acetylated tubulin as a ciliated cell marker (green). Graphs show quantification of dots per cell type, per genotype, for each probe. Scale bar=30 μm. (n=3 mice per group). (A) Secretory vs ciliated (18a^F/F^), P=0.03; secretory (18a^F/F^) vs secretory (18a^Δ/Δ^), P=0.0038; ciliated (18a^F/F^) vs ciliated (18a^Δ/Δ^), P=0.0254. (B) Secretory vs ciliated (18b^F/F^), P=0.0012; secretory (18b^F/F^) vs secretory (18b^Δ/Δ^), P=0.0002; ciliated (18b^F/F^) vs ciliated (18b^Δ/Δ^), P=0.0253. (C) Secretory (18c^F/F^) vs secretory (18c^Δ/Δ^), P=0.0017; ciliated (18c^F/F^) vs ciliated (18c^Δ/Δ^), P=0.0004, Student’s two-tailed *t* test. #, P<0.05 between cell types; *, P<0.05 within cell type.

### Munc18b predominates in stimulated mucin secretion

Our previous study using heterozygous mutant mice indicated that Munc18b has a major role in stimulated mucin secretion (23). To comprehensively analyze Munc18 function in stimulated secretion, mucin production was first increased (mucous metaplasia) in all mutant mice using ovalbumin sensitization and challenge to induce allergic inflammation, and then mice were stimulated with the secretagogue ATP to induce secretion acutely (Figure 3, A). None of the Munc18 floxed mice showed a phenotype in mucous metaplasia or in mucin secretion in this or any subsequent experiments. All three Munc18 single conditional deletants had mucin content similar to WT mice after the ovalbumin challenge, but the Munc18a/b double deletant had significantly higher mucin content (Figure 3, A and B), suggesting a defect in baseline secretion that results in mucin accumulation (see below). After ATP exposure, the predominant role of Munc18b was confirmed because the conditional deletant secreted only ~26% of intracellular mucin (mean value) compared to Munc18b floxed mice that secreted ~60% or WT mice that secreted ~52% (Figure 3, C). Munc18a and Munc18c deletants secreted as efficiently as their cognate floxed mice or WT mice, while the Munc18a/b double deletant secreted ~31%, comparable to the Munc18b single deletant (Figure 3, C).

**Figure 3.**
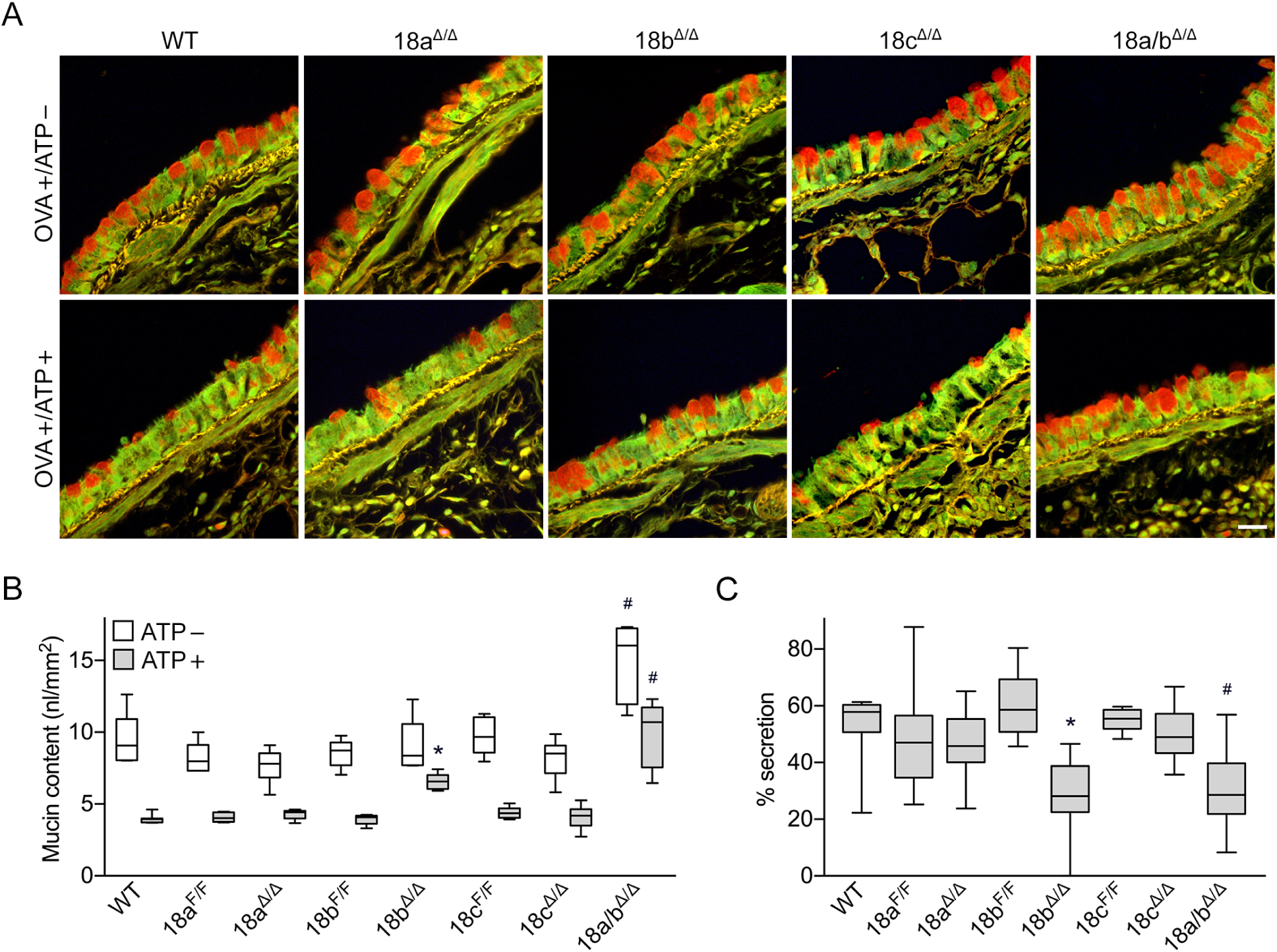
Stimulated mucin secretion measured by residual intracellular mucin content. **(A)** High-magnification views of representative fields of PAFS-stained bronchial airways from mice sensitized and challenged with ovalbumin to increase mucin production (OVA+/ATP-, top row), then exposed to aerosolized 100 mM ATP to stimulate mucin secretion (OVA+/ATP+, bottom row). Scale bar=20 μm. **(B)** Quantification of the volume density (expressed as nl mucin per mm^2^ basement membrane) of intracellular mucin in mice with or without ATP stimulation as in (A) (representative experiment of three separate experiments with all genotypes, n=5-8 mice per group). 18a/b^Δ/Δ^ (ATP-) vs WT (ATP-), P<0.0001, vs 18a^F/F^ (ATP-), P<0.0001, vs 18b^F/F^ (ATP-), P<0.0001, Tukey test. 18b^Δ/Δ^ (ATP+) vs 18b^F/F^ (ATP+), P<0.0001; 18a/b^Δ/Δ^ (ATP+) vs WT (ATP+), P<0.0001, vs 18a^F/F^ (ATP+), P=0.0001, vs 18b^F/F^ (ATP+), P<0.0001, Student’s two-tailed *t* test. **(C)** The percentage of mucin released for each genotype (three independent experiments like those in (B), combined). 18b^Δ/Δ^ vs 18b^F/F^, P<0.0001; 18a/b^Δ/Δ^ vs WT, P=0.001, vs 18a^F/F^, P<0.0001, vs 18b^F/F^, P<0.0001, Mann-Whitney test. Box plots, line=median; box=25^th^-75^th^ percentile; whiskers=5^th^-95^th^ percentile for this and all subsequent figures. *, P<0.05 vs floxed littermate; #, P<0.05 vs WT.

### Munc18a predominates in baseline mucin secretion

No defect in baseline mucin secretion had been observed in Munc18b heterozygous mutant mice (23). This could be due either to the lack of a role of Munc18b or to the lack of an obvious phenotype with just a 50% reduction in protein expression. To further study baseline mucin secretion, all Munc18 airway deletants were examined.

In naïve (uninflamed) WT mice, the rate of mucin secretion closely matches the rate of mucin production such that intracellular mucin does not accumulate and is not visible by PAFS histochemical staining. Hence, a defect in baseline mucin secretion can be detected as spontaneous mucin accumulation (15, 30). Munc18a deletant mice showed significant spontaneous mucin accumulation, indicating a role of Munc18a in baseline secretion (Figure 4, A and B). Munc18b and Munc18c deletants showed no mucin accumulation. However, Munc18a/b double deletant mice showed a higher level of mucin accumulation than Munc18a single deletant mice, indicating an additive effect of Munc18a and Munc18b in baseline mucin secretion. Spontaneous mucin accumulation was further analyzed by quantitative immunoblotting for Muc5b, which is expressed in the airways of naïve mice (4, 8, 15, 31). This confirmed significant mucin accumulation in Munc18a deletant mice, a trend towards a small increase in Munc18b deletant mice (P=0.09), and an additive effect in Munc18a/b double deletant mice (Figure 4, C; Figure S4). Muc5ac is not expressed significantly in the airways of naïve mice (4, 8, 31), and was not detected in immunoblots of the lungs of any naïve Munc18 deletant mice (not shown). To rule out the possibility that spontaneous mucin accumulation was due to an increase in mucin expression resulting from an inflammatory response in any of the deletants, mRNA expression was analyzed by qRT-PCR in Munc18a and Munc18 single deletant mice and Munc18a/b double deletants. There was no significant increase in expression of *Muc5ac* or *Muc5b* in any of the conditional deletants (Figure S6, A).

**Figure 4.**
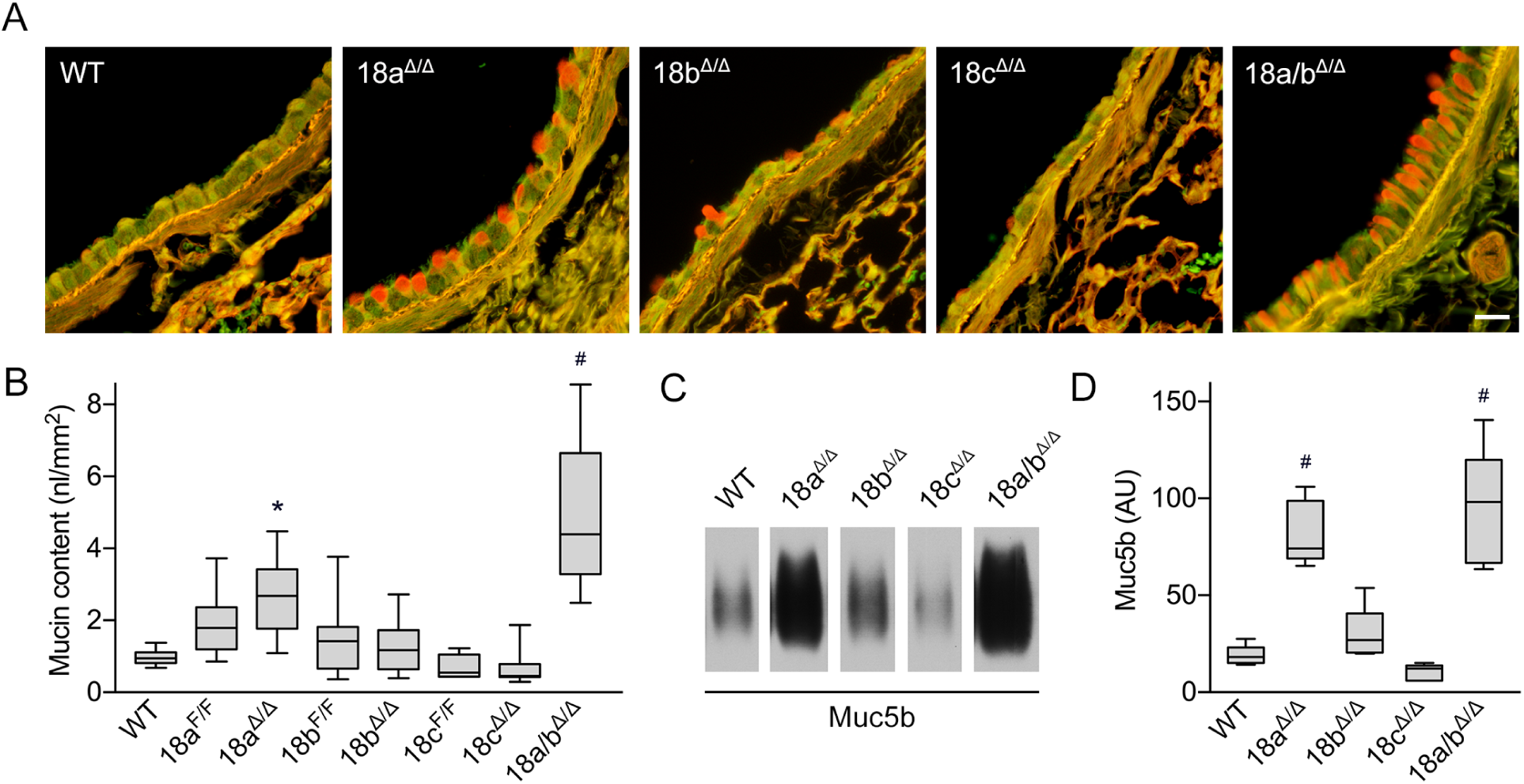
Baseline mucin secretion measured by spontaneous intracellular mucin accumulation. **(A)** High-magnification views of representative fields of PAFS-stained bronchial airways from naïve mice. Scale bar=20 μm. **(B)** Quantification of the volume density of spontaneous intracellular mucin accumulation (n=5-18 mice per group, three independent experiments combined). 18a^Δ/Δ^ vs 18a^F/F^, P=0.0243; 18a/b^Δ/Δ^ vs WT, P<0.0001, vs 18a^Δ/Δ^, P=0.0008, Student’s two-tailed *t* test. **(C)** Representative immunoblot of 50 μg of whole lung lysates from naïve mice probed for Muc5b. **(D)** Densitometric analysis of immunoblot shown in (C) derived from standard curve (Fig. S4) (AU, arbitrary units) (n=5-7 mice per group). 18a^Δ/Δ^ vs WT, P=0.0079; 18a/b^Δ/Δ^ vs WT, P=0.0025, Mann-Whitney test. *, P<0.05 vs floxed littermate; #, P<0.05 vs WT.

To examine granule morphology by electron microscopy and stereological analysis, mice were lightly stimulated with IL-13 so that mucin granules were readily visible in WT mice (Figure S5, A). Munc18a/b double deletant mice showed an increase in surface-to-volume density of granules (Sv, Figure S5, B), indicating smaller granules with a lower ratio of surface area to volume. However, there was no change in volume density of granules per volume density of cells (Vv, Figure S5, C). Together, these findings indicate that the cells from Munc18a/b double deletant mice had an increased number of smaller granules. In addition, mucin granules in Munc18a/b double deletant mice were also found to be more electron-dense (Figure S5, D).

### Deletion of Munc18a or Munc18b does not impair mucociliary clearance

The absence of an increase in mucin transcripts in the Munc18 airway deletants suggested that mucociliary clearance function was preserved, preventing lung infection and inflammation that might induce mucin gene upregulation (Figure S6, A). To further test these inferences, several additional studies were performed. First, lung lavage fluid was obtained for measurement of leukocytes, which is a sensitive indicator of inflammatory status. There was no difference in total cell number or fractional representation of any leukocyte subset in either of the Munc18a or Munc18b single deletant mice compared to WT or floxed littermate mice (Figure S6, B). However, the double deletant mice showed a small but significant increase in neutrophil number. Next, polystyrene beads were instilled into the lungs to measure their clearance by mucociliary transport. There was no difference in the fraction of beads cleared by Munc18a or Munc18b deletant mice compared to WT (Figure S6, C). Last, the lung microbiome was interrogated by qPCR and sequencing of 16S ribosomal RNA. There was no significant difference in the quantity (Figure S6, D) or composition (Figure S6, E) of bacteria present in the lungs of Munc18 single or double deletant mice compared to WT or floxed littermate mice.

### Deletion of Munc18b protects mice against airway obstruction in a model of allergic asthma

Increased mucin production followed by stimulated secretion causes airway lumenal mucus occlusion and airflow obstruction in asthma (1, 12). We hypothesized that deletion of Munc18b in airway secretory cells would protect against this pathophysiology. To test this, we first performed a pilot study in a WT mouse with IL-13 instilled intrapharyngeally to induce mucous metaplasia and then exposed to a methacholine aerosol to stimulate mucin secretion. We measured the occlusion of airways throughout the lungs at 500 μm intervals as a fraction of cross-sectional airway area, and found that the right caudal lobe had the highest fractional occlusion (Figure S7, A). We next compared fractional occlusion in the right caudal lobe between Munc18b floxed and deletant mice and found a significant reduction (~50%, mean values) in the deletant mice (Figure S7, B). We then performed a definitive study comparing the sum of the area of lumenal mucus in the right caudal lobe at 1 mm intervals together with measurement of lung mechanics (Figure 5). Munc18b deletant mice showed a ~62% reduction (mean values) in lumenal mucus area compared to Munc18b floxed mice, similar to the ~37% reduction in Muc5ac knockout mice, whereas Munc18a deletant mice showed no reduction (Figure 5, B).

**Figure 5.**
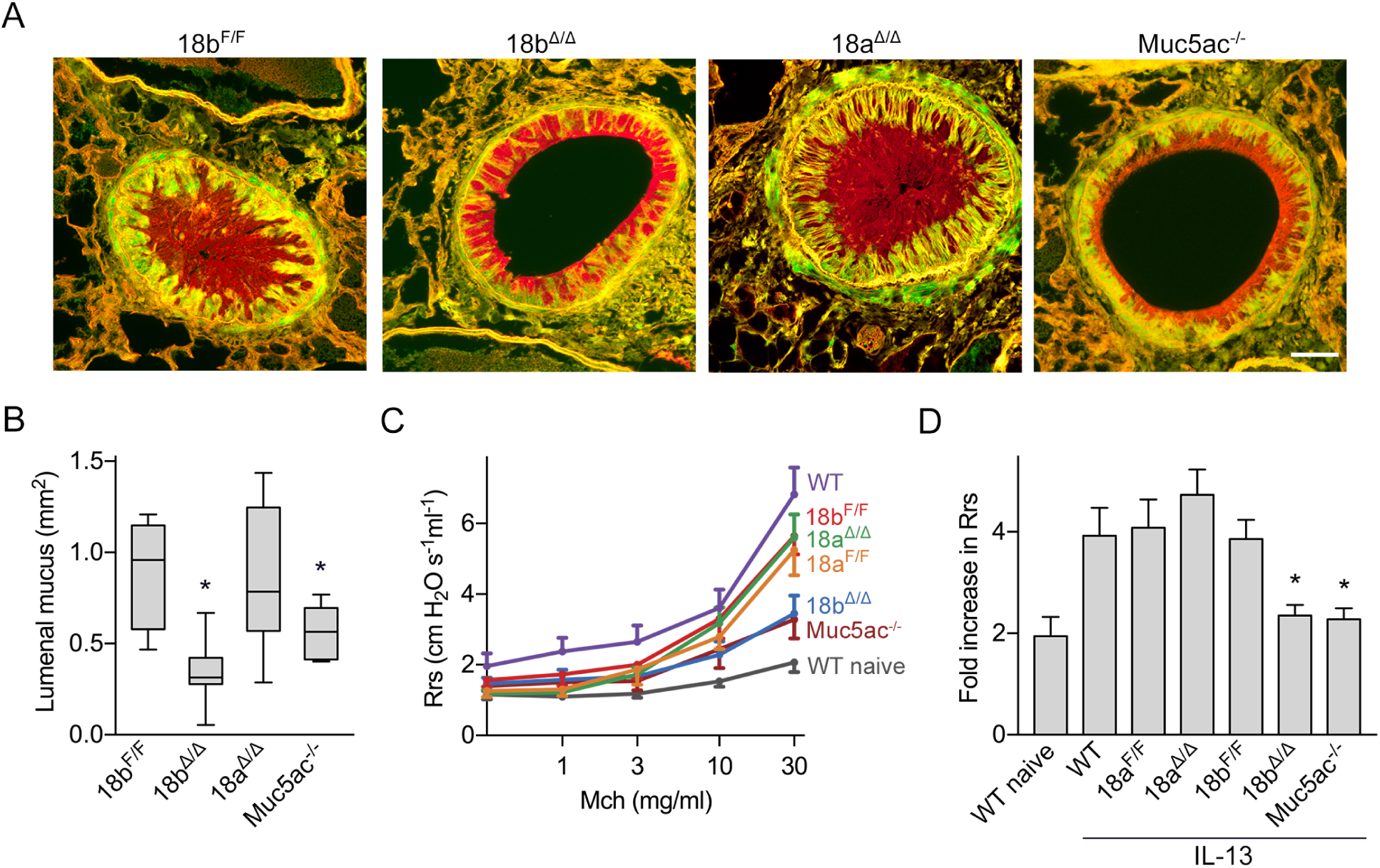
Airway mucus occlusion and airway hyperreactivity of Munc18b conditional deletant mice in an allergic asthma model. **(A)** Representative cross-sections of airways fixed with methacarn to preserve mucus volume and stained with PAFS. Mice were treated with IL-13 to induce mucin production and then stimulated with aerosolized 150 mM methacholine (Mch) to stimulate mucin secretion and smooth muscle contraction. Scale bar=50 μm. **(B)** Cross-sectional area of lumenal mucus in the right caudal lobe measured at 1000 μm intervals. (n=5-9 mice per group, representative experiment, >100 airways per group quantified). 18b^Δ/Δ^ vs 18b^F/F^, P=0.0002; Muc5ac^-/-^ vs 18b^F/F^, P=0.0378, Student’s two-tailed *t* test. **(C)** Total respiratory system resistance (Rrs) at increasing doses of nebulized Mch in mice treated with or without IL-13 (n=6-14 per group, three independent experiments combined). Line, mean; error bar, SEM. **(D)** Fold-change Rrs at the highest dose of nebulized Mch (30 mg/ml) from (C). Each genotype is normalized to its own baseline measured with nebulized saline. 18b^Δ/Δ^ vs 18b^F/F^, P=0.0018; Muc5ac^-/-^ vs 18b^F/F^, P=0.0044, Student’s two-tailed *t* test. *, P<0.05 vs floxed littermate. Bar, mean; error bar, SEM.

Exposure to methacholine induces resistance to airflow due to a combination of smooth muscle contraction (bronchoconstriction) and mucus obstruction (12). An augmented response to methacholine (airway hyperresponsiveness) is a sensitive indicator of asthmatic airway dysfunction. WT mice with mucous metaplasia induced by IL-13 and exposed to increasing concentrations of aerosolized methacholine showed increased respiratory system resistance compared to naïve WT mice (Figure 5, C and D). Munc18a^F/F^, Munc18b^F/F^, and Munc18a^Δ/Δ^ mice with mucous metaplasia were similar to WT mice, but Munc1 8b^Δ/Δ^ and Muc5ac^-/-^ mice with mucous metaplasia were highly protected from airway hyperresponsiveness to methacholine (Figure 5, C and D).

### Deletion of Munc18b protects mice against airway mucus occlusion and parenchymal emphysema in a model of cystic fibrosis

In cystic fibrosis, an inherited defect in transepithelial anion and water transport causes the formation of mucus that is excessively concentrated, viscoelastic and adhesive (32–34). Mucus accumulates because its excessive viscoelasticity impedes clearance by ciliary beating and its adhesivity results in the formation of airway mucus plaques. These in turn, lead to infection and inflammation that cause progressive lung disease. In mice, deletion of the anion transporter, CFTR, does not result in lung disease because of the presence of alternative mechanisms of anion transport, but transgenic overexpression of the beta subunit of the epithelial Na^+^ channel (ENaC) results in concentrated mucus leading to lung disease that resembles human cystic fibrosis (35).

To test whether stimulated mucin secretion contributes to pathophysiology in this model, we crossed Munc18b deletant mice with βENaC-Tg mice (Figure 6, A). Mucus occlusion in βENaC-Tg mice was reduced by ~66% (mean values) by deletion of Munc18b in the airway (Figure 6, B). Emphysema measured as equivalent mean diameter (D_2_) was increased ~86% in βENaC-Tg-Munc18b^F/F^ mice compared to Munc18^F/F^ mice as previously described for the βENaC-Tg (35) (Figure 6, C). This increase was attenuated by ~22% by deletion of Munc18b in the airway (Figure 6, C). However, lung neutrophilic and eosinophilic inflammation present in βENaC-Tg-Munc18b^F/F^ was not reduced by Munc18b deletion (Figure 6, D).

**Figure 6.**
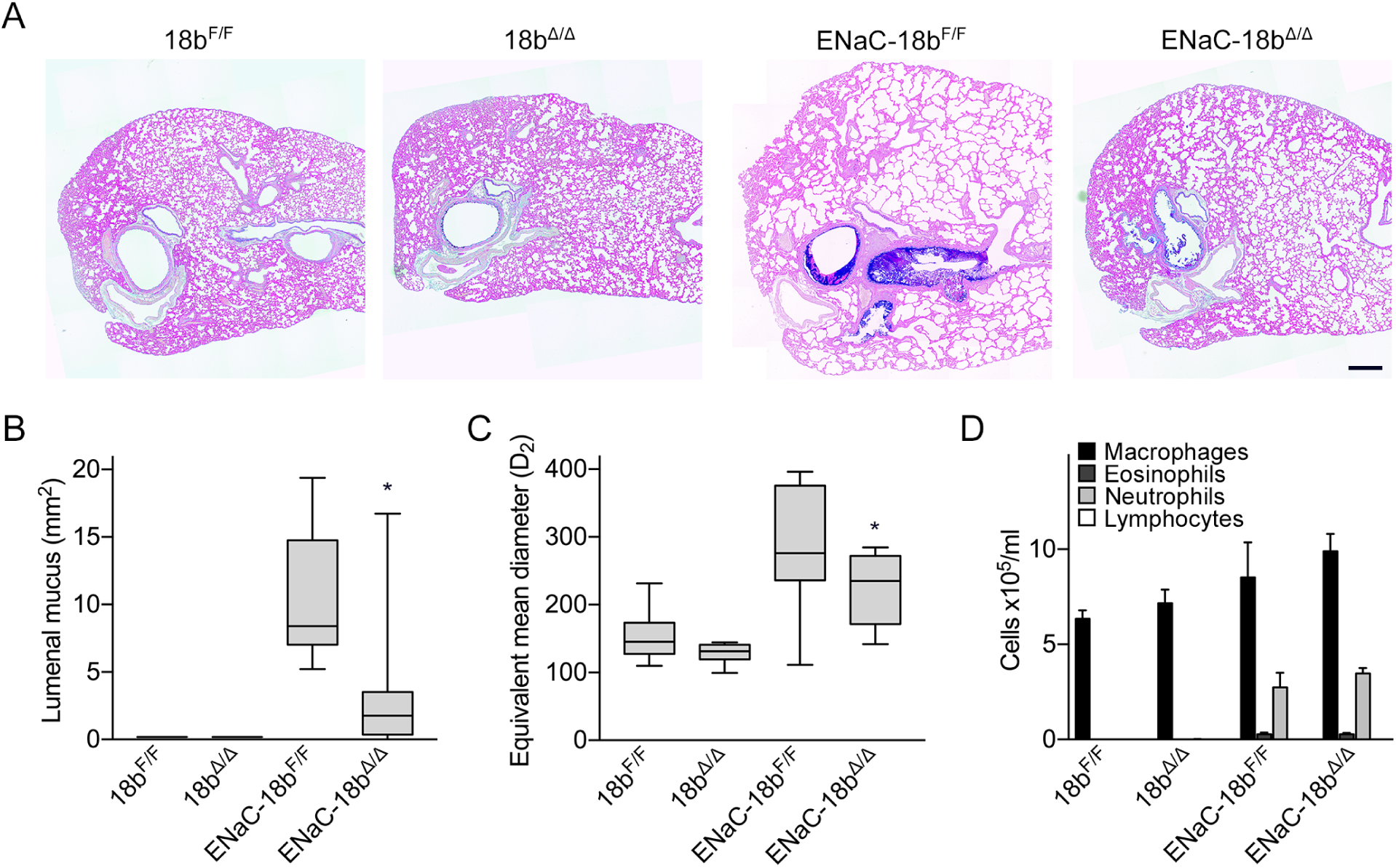
Airway mucus occlusion and emphysema of Munc18b conditional deletant mice in a cystic fibrosis-like model. Munc18b^F/F^ and Munc18b^Δ/Δ^ mice were crossed or not to β-ENaC-overexpressing transgenic (ENaC) mice. **(A)** Representative transverse left lung sections stained with AB-PAS. Scale bar=300 μm. **(B)** Cross-sectional area of lumenal mucus was quantified. (n=8-16 mice per group). ENaC-18b^Δ/Δ^ vs ENaC-18b^F/F^, P=0.0013, Mann-Whitney test. **(C)** Emphysema was assessed as the equivalent mean diameter, D_2_. (n=8-16 mice per group). ENaC-18b^Δ/Δ^ vs ENaC-18b^F/F^, P=0.0009, Student’s two-tailed *t* test. **(D)** Total inflammatory cell numbers from lung lavage fluid (n=8-16 mice per group). *, P<0.05 vs floxed littermate.

## Discussion

Here, we have performed a comprehensive analysis of the function of Munc18 proteins in a polarized epithelial cell specialized for apically directed regulated secretion. Our central finding is that baseline and stimulated secretion are predominantly mediated by different Munc18 proteins, with Munc18a having the major role in baseline secretion, Munc18b having the major role in stimulated secretion, and Munc18c having no apparent role. This finding has implications for understanding the cell biology of regulated secretion in polarized cells and for manipulating the exocytic machinery of the airway epithelium therapeutically to alleviate mucus dysfunction.

Regarding cell biology, Munc18b has been described previously by us and others as mediating apical secretion in polarized epithelial cells (23, 36, 37). Munc18a has been studied primarily for its role in synaptic vesicle release from neurons (24, 38, 39), but other regulated exocytic systems where Munc18a has been reported to function include vascular endothelial cells (40) and acrosomal exocytosis in spermatozoa (41). In airway epithelium, Munc18a was reported to modulate the conductance of the apical anion channel CFTR (42), but a role in vesicular transport was not described. Munc18a and Munc18b have been reported to cooperate in regulated exocytosis of insulin-containing granules of pancreatic islet cells (43, 44) and cytolytic granules of natural killer cells (45) in response to different signaling pathways. Prior reports that Munc18a and Munc18b both participate in mast cell degranulation appear to have been in error, with Munc18b having an exclusive role as we recently described (26). Thus, the current work is the first report, of which we are aware, of two different Munc18 proteins mediating different rates of secretion in response to the same signaling pathway.

Munc18c has been proposed to mediate basolateral secretion in polarized epithelial cells (46, 47), consistent with the lack of effect of Munc18c deletion in apical regulated secretion in our system (Figures 3 and 4). A role for Munc18c in stimulated exocytosis has been described in non-polarized cells, such as translocation of glucose transporters in adipocytes (48), but not in in polarized epithelia. The cellular lethality induced by simultaneous deletion of both Munc18b and Munc18c in airway epithelial cells (Figure S3) suggests that the predominant function of Munc18c is in constitutive secretion because that is an essential cellular function. We hypothesize that Munc18c function is rescued in the single airway deletant mouse by ectopic function of Munc18b. This hypothesis is consistent with the cellular viability of airway epithelial cells in double Munc18a/b mice (Figures 3 and 4) because regulated secretion is not a cell-autonomous essential function. Whether Munc18c mediates constitutive secretion at both the apical and basolateral surfaces in airway epithelial cells or functions exclusively at the basolateral surface is not known.

Since Munc18 proteins partner with specific Syntaxins t-SNAREs (23), our findings here suggest that the exocytic SM-SNARE machinery is mostly different between baseline and stimulated mucin secretion. This inference is supported by the role of the v-SNARE VAMP8 predominantly in stimulated secretion (21), even though SNAP23 functions in both processes (20). What might be the adaptive value of utilizing different exocytic machines for baseline and stimulated secretion rather than utilizing a single machine capable of running at different rates? In view of the differing roles of baseline and stimulated mucin secretion in mucociliary clearance and airway occlusion (12), and the differing roles of Muc5ac and Muc5b in helminthic and microbial defense (4, 10), several plausible possibilities exist. These include different exocytic machines acting on small immature granules in baseline secretion to minimize the chance of airway occlusion, and on large mature granules in stimulated secretion to maximize the chance of occlusion. This might occur by exchanging VAMP proteins during granule maturation, by analogy with Rab conversion during progression from early to late endosomes (49). Another possibility is that Muc5ac and Muc5b are packaged separately during exit from the trans-Golgi network (50), with different exocytic machines acting on secretory granules containing either secreted mucin. This last possibility is supported by the apparent segregation of MUC5AC and MUC5B extracellularly and intracellularly in human asthmatic airways (51). Further studies colocalizing vesicular components of the exocytic machinery such as VAMPs and Synaptotagmins with different secreted mucins will be required to resolve these questions. The fact that secretory granules visualized by EM in Munc18a/b double deletant mice are smaller and denser than granules in WT mice (Figure S5) suggests that these SM proteins also mediate homotypic fusion between granules during post-Golgi maturation, and that mucins decondense to some degree during granule maturation.

Importantly, the existence of two different exocytic machines in airway secretory cells affords the possibility to molecularly target the pathologic consequences of stimulated mucin secretion without compromising the critical protective clearance function of baseline mucin secretion. While the fundamental pathophysiologic processes in allergic asthma and cystic fibrosis are mucin hyperproduction and impaired anion transport, respectively, stimulated mucin secretion contributes to airway occlusion in both diseases as indicated by our prior (12) and current studies in mouse models of allergic asthma (Figure 5) and cystic fibrosis (Figure 6). To fully appreciate the contribution of stimulated secretion to airway mucus occlusion, it is important to recognize several features of mucin biology. First, the production of mucins, particularly of Muc5ac, can be greatly increased by inflammatory signaling, resulting in the filling of airway epithelial secretory cells with large amounts of mucin contained within secretory granules (Figure 3). If secretion is not stimulated, the stored mucins are slowly released and cause only minimal mucus occlusion (12). Second, stimulated secretion can result in the explosive exocytic release of mucin granule contents within seconds (18), and the massive extracellular swelling of the released mucins by absorbing several hundred-fold their mass of water occurs in less than a second (3, 52). Thus, the area of the airway cross-section occupied by intracellular mucin is only a small fraction of that occupied by fully hydrated extracellular mucus, and the extracellular expansion of mucin volume within such a short time frame are key determinants of pathophysiology. In large proximal airways, rapid release of stored mucins from surface epithelial cells is unlikely to completely occlude the airway lumen, and may have adaptive value in promoting the trapping of particles and pathogens for clearance, as described for submucosal glands (53, 54). However, in small distal airways, rapid massive mucin release can overwhelm clearance mechanisms, resulting in airway lumenal occlusion (12) (Figures 5 and 6).

The adaptive value of small airway occlusion in the stimulated secretion of metaplastic epithelial cells may be to trap helminths migrating through lungs (11), a hypothesis being tested by us and others. In allergic asthma, IL-13 plays a central role in increased mucin production, as it does in helminth infection, and rapid secretion can be induced by acute inflammatory mediators such as acetylcholine and histamine (11, 12). In cystic fibrosis, increased mucin production is more modest (32), and the role of secretagogues in small distal airways is less well studied. However, the stimulation of secretion from submucosal glands is critical to mucus dysfunction in a pig model of cystic fibrosis (32), and the stimulation of mucin secretion from surface epithelial cell by acetylcholine from neurons or ATP from leukocytes or epithelial cells could contribute to mucus occlusion in small airways.

Mucus occlusion was significantly reduced by deletion of Munc18b in airway epithelial cells in both of the mouse models of airway disease we tested. In the allergic asthma model, reduction of mucus occlusion was shown to result in improvement of lung mechanics (Figure 5, C and D). In the cystic fibrosis model, while the emphysema that occurs secondary to mucus occlusion was also mitigated (Figure 6, A and C), neutrophilic and eosinophilic inflammation was not reduced (Figure 6, D), similar to what occurred upon treatment with aerosolized hypertonic saline solution in that model (55). Deletion of Munc18b in airway epithelium did not result in any abnormalities of lung structure, particle clearance, inflammation, or bacterial infection (Figure S6). Therefore, targeting the stimulated exocytic machine with small molecules or RNA silencing technologies in human subjects might be free of intrinsic adverse consequences.

Whether the trapping of helminths during migration through the lungs might be impaired by Munc18b deletion is an area of our active investigation, but helminth infestation is not a common problem in developed countries.

## Methods

### Mice

C57BL/6J (catalog no. 000664), C57BL/6-Tg(Zp3-cre)93Knw/J (catalog no. 003651), Gt(ROSA)26Sor^tm4(ACTB-tdTomato,-EGFP)Luo^/J (catalog no. 007576) and B6N.129S6(Cg)-Scgb1a1^tm1(cre/ERT)Blh/^/J (catalog no. 016225) mice were purchased from The Jackson Laboratory. We obtained Munc18a conditional deletant mice from Dr. Matthijs Verhage (U. Amsterdam) (27), Munc18c conditional deletant mice from Dr. Roberto Adachi (U. Texas MD Anderson Cancer Center) (26) and CCSP^iCre^ (Scgb1a1^tm1(icre)Fjd^) mice from Dr. Francisco DeMayo (NIEHS) (28). β-ENaC-C57BL/6-Tg mice (35) were crossed to Munc18b conditional deletant mice at the University of North Carolina at Chapel Hill.

In the Munc18a conditional deletant, exon 2 of the gene is flanked by two loxP sites and Cre recombination induces a frameshift resulting in a nonsense codon and absence of protein (27). In the Munc18c conditional deletant, exon 1 is flanked by two loxP sites and Cre recombination removes the start codon (26). To generate Munc18b conditional deletant mice, we built a targeting vector to insert two loxP sites to flank exon 1 (Figure S1) by homologous recombination. Upstream of exon 1, the zeocin and puromycin resistance genes flanked by two loxP sites (M2) were inserted, and downstream of exon 1, the phosphoglucokinase promoter-neomycin resistance gene (PGK-neo) resistance gene flanked by two Flp recognition target (FRT) sites was inserted. The herpes simplex virus thymidine kinase gene was introduced outside the homology arms for a negative selection marker. This vector was electroporated into 129S6:B6 embryonic stem cells; after 24 h cells were selected using (1-(2-deoxy-2-fluoro-1-D-arabinofuranosyl)-5-iodouracil), puromycin and G418. Of 84 surviving clones, one correctly targeted clone was chosen for subsequent manipulation. The puromycin resistance gene was removed in vitro by electroporation of a circular CMV-Cre plasmid into the positive clone. Cells were plated at low density (2.5 × 10^5^ to 2.0 × 10^4^ cells/ml), and individual clones were transferred to 96-well plates. Duplicate plates were prepared, and one plate selected on puromycin. Of the 19 subclones that did not survive puromycin selection, one clone was shown to have correct targeting to remove the puromycin resistance gene by PCR. This clone was used for injection into B6(Cg)-Tyrc-2J/J blastocysts. The 8 chimeric males generated were crossed to B6(Cg)-Tyrc-Gt(ROSA)26Sor^tm1(FLP1)Dym^/RainJ (The Jackson Laboratory, catalog no. 009086) to remove PGK-Neo and establish our floxed line. We then crossed the Munc18b floxed mouse with C57BL/6-Tg(Zp3-cre)93Knw/J (catalog no. 003651) to generate full animal KOs (Figure 1, B) or with CCSP^iCre^ mice for specific deletion in the airway epithelium. All our lines were crossed with C57BL/6J mice for 10 generations.

Genotypes of Munc18b mutant mice were determined by PCR of genomic DNA with primers #7 (AAGGCGGTGGTAGGGAAAGT) and #64 (CAGTTGGTCAAATTCAAGTGCTC) to differentiate between the conditional (F; 1075 bp) and WT (+; 931 bp) alleles. Munc18a and Munc18c were genotyped as previously described (26). The presence of Zp3-cre was determined by PCR with primers ZpCre 5’ (GCGGTCTGGCAGTAAAAACTTC); ZpCre 3’ (GTGAAACAGCATTGCTGTCACTT); IntControl 5’ (CTAGGCCACAGAATTGAATTGAAAGATCT); IntControl 3’ (GTAGGTGGAAATTCTAGCATCATCC), that give a 324 bp internal control band and a 100 bp band from the transgene. The presence of conditional CCSP^iCre^ was determined by PCR with primers CC10-iCreR (GAGATGTCCTTCACTCTGATTC); CC10-iCreF (TCTGATGAAGTCAGGAAGAACC); FJD13 (TGCCAGAGATTGTTCTAGAAAACAA) and FJD14 (GGCACAATGATGTTAATGACGTAAA), that give a 1 kbp internal control band and a 0.5 kbp band from the transgene. Genotyping for the β-ENaC-Tg mice was performed as previously described (35).

For CreER induction, five doses of 6 mg of tamoxifen (T5648, MilliporeSigma) dissolved in corn oil (C8267, MilliporeSigma) were injected into adult mice every other day intraperitoneally (i.p.). Mice were harvested two days after the last dose. Mice of both sexes were used in all experiments, ranging from 6-32 weeks of age.

### Immunohistochemistry

Lungs were inflated and fixed with 10% neutral buffered formalin (NBF) for 24 h at 4°C and then embedded in paraffin. Lung sections were cut into 5-μm transverse sections, deparaffinized, exposed for 10 min to 3% H_2_O_2_ in 90% methanol and then heated for 10 min in 10 mM sodium citrate, pH 6.0, for antigen retrieval. Tissue sections were blocked with 5% donkey serum (017-000-121, Jackson ImmunoResearch) for 1 h at room temperature and then incubated with goat anti-CCSP (a gift from Dr. Barry Stripp, Cedars-Sinai, 1:2000) diluted in blocking solution at 4°C overnight. Secondary antibody-horseradish peroxidase (HRP)-labeled donkey anti-goat (705-035-003, Jackson ImmunoResearch, 1:500) was incubated for 1 h at room temperature. Tissue sections were then washed with PBS, dehydrated and mounted with VectaMount (Vector Laboratories).

### In situ hybridization and immunofluorescence

*In situ* RNA detection was performed using the RNAscope detection kit (Advanced Cell Diagnostics, Hayward, CA) according to the manufacturer’s instructions. Briefly, lungs were inflated and fixed with 10% NBF for 30 h at room temperature and then embedded in paraffin. Tissue blocks were cut into 5-μm sections, deparaffinized, and pretreated with heat and protease before hybridization with the target oligonucleotide probes: murine Munc18a (Probe-Mm-Stxbp1, 521961), Munc18b (Probe-Mm-Stxbp2, 536201) and Munc18c (Probe-Mm-Stxbp3, 536191). The positive control probe was Ppib (peptidylprolyl isomerase B, 313911) and the negative control probe was DapB (4-hydroxy-tetrahydrodipicolinate reductase from *Bacillus subtilis*, 310043). Preamplifier, amplifier and fluorescent-labeled oligonucleotides were then hybrized sequentially. RNAscope signal was imaged at the axial bronchus, between lateral branches 1 and 2 (30). Images were acquired using a confocal microscope (A1plus, Nikon) with a 40× NA 1.3 objective lens. For analysis, the number of dots per secretory or ciliated cell were counted using ImageJ (56). Since dot intensity represents transcript amount, brighter and/or bigger dots were counted twice. More than 70% of the dots were scored as singlets. Scoring all the doublets as singlets can only result in a 4% error.

Immunofluorescence was performed using antibodies against goat CCSP (1:2000) and mouse acetylated tubulin (T6793 MilliporeSigma, 1:2000). Secondary antibodies were species-specific donkey anti-IgG coupled with Alexa Fluor 488 (A21202) and Alexa Fluor 647 (A21447) (1:1000, Invitrogen).

### Airway epithelial mucin content by PAFS staining and image analysis

Mucous metaplasia was induced in the airways of mice by sensitizing mice to ovalbumin (OVA) (20 μg OVA Grade V, 2.25 mg alum in 0.9% saline, pH 7.4; MilliporeSigma) administered by i.p. injection once weekly, for 3 weeks. Sensitized mice were exposed for 20 min to an aerosol challenge of 2.5% (wt/vol) OVA in 0.9% saline supplemented with 0.02% (vol/vol) antifoam A silicon polymer (MilliporeSigma) daily for five days via an Aerotech II compressed gas nebulizer (Biodex, New York) in the presence of room air supplemented with 5% CO_2_. Three days after the last OVA aerosol exposure, half of the mice in each group were exposed for 10 min to an aerosol of 100 mM ATP (MilliporeSigma) in 0.9% NaCl to induce mucin secretion, then sacrificed after 20 min. Lungs were harvested, inflated and fixed in 10% NBF, embedded in paraffin and sectioned into a single transverse 5 μm cut of the axial airway of the left lung, between the lateral branches 1 and 2 (30). Sections were deparaffinized, rehydrated, and then stained with periodic acid fluorescent Schiff reagent (PAFS). Images were acquired using an upright microscope (Olympus BX 60) with a 40× NA 0.75 objective lens and intracellular mucin was measured around all the circumferential section of the axial bronchus using ImagePro (Media Cybernetics, Bethesda, MD). Data are presented as the epithelial mucin volume density, signifying the measured volume of mucin overlying a unit area of epithelial basal lamina, derived as described (57). Images were analyzed by investigators blinded to mouse genotype and treatment.

### Agarose gel Western blot for mucin

Lungs were perfused by intracardiac injection of 2 ml PBS until blanched, then homogenized in 1 ml of 6 M guanidinium buffer containing protease inhibitors and incubated for two days at 4°C. Lysates were centrifuged at 19,000 g and the supernatants were then dialyzed overnight at 4°C against PBS in Slide-A-Lyzer 2K MWCO 3 ml cassettes (Thermo Fisher Scientific). Protein concentrations were determined using a bicinchoninic acid protein assay kit (Thermo Fisher Scientific) and then the samples were incubated with 20 Kunitz DNase (LS002139, Worthington) for 15 min at 37°C and reduced with 10 mM DTT (MilliporeSigma) for 10 min at 95°C in loading buffer (5% glycerol, 0.1% SDS, 0.0025% bromophenol blue, 0.6 M urea, 10 mM Tris-HCl, 0.5 mM EDTA, pH 8). Samples were electrophoresed through a 0.8% agarose/0.1% SDS hydrogel, and the gel was then soaked in 10 mM DTT for 20 min, then the samples were transferred by vacuum onto a PVDF membrane. Membranes were washed in PBS and blocked with 5% nonfat milk in PBS/0.05% Tween before probing with lectin UEA-1 conjugated to HRP (1:1000, L8146, MilliporeSigma) in blocking solution to detect fucosylated Muc5ac (30), or with monoclonal antibody MDA-3E1 raised by us against peptide TTCQPQCQWTKWIDVDYPSS in blocking solution (1:1500) to detect Muc5b (20). Secondary antibody for Muc5b was goat anti-mouse conjugated to HRP (1:5000, Thermo Fisher Scientific), and the chemiluminescence signal was detected with ECL Western Blotting Substrate (Thermo Fisher Scientific). Relative protein amounts were determined for each sample using ImageJ (NIH) against a standard curve run for each gel (Figure S4).

### Electron microscopy and stereology

Slight mucous metaplasia was induced in mice with one dose of 0.2 μg of IL-13 instilled intrapharyngeally on day 1 of the experiment in order to achieve optimal visualization of mucin granules. Mice were then anesthetized and sacrificed on day 5. Lungs were excised, fixed in 2.5% glutaraldehyde in 0.1 M sodium cacodylate buffer (pH 7.2), containing 20 mM calcium chloride for 2 h, followed by a 1 h post-fixation in buffered 1% osmium tetroxide. The fixed left lung was then sectioned into a single transverse cut of the axial airway between lateral branches 1 and 2 and embedded in Embed 812 epoxy resin (14120 EMS). Sections of 100 nm thickness were stained with uranyl acetate and lead citrate and were viewed at 8200× magnification in a Tecnai 12 transmission electron microscopy. Secretory cells were randomly selected and imaged, and at least 19 images were used per mouse. Vv and Sv were obtained with randomly placed dot and line grids (line pairs, 64 tiles) on the cell profiles (58). To measure the relative electron lucency of mucin granules, a cycloid stereological grid with circles of 30 pixels in diameter was randomly superimposed on the images. At least 5 circles that fell on nuclear heterochromatin and electron-lucent extracellular space were selected, and their grayscale value (0–255) was recorded. These values were used to set a linear scale for each image. Then at least 30 random circles that fell on granules were recorded using that scale. These data were recorded and the appropriate bin range and histogram distribution was calculated.

### Mucin transcript quantitative RT-PCR analysis

Total RNA was extracted from whole lung (RNeasy mini kit; Qiagen) and reverse-transcribed (iSCRIPT, Bio-Rad). Quantitative PCR was carried out for each cDNA sample in triplicates with qPCR master mix (Quanta Biosciences) and 6-carboxyfluorescein-labeled probes for Muc5b (Mm00466391_m1), Muc5ac (Mm01276725_g1) and β-actin (Mm02619580_g1; all from Thermo Fisher Scientific) on a ViiA 7 RT PCR System (Applied Biosystems). Results were expressed as ΔCt (normalized for β-actin) (59).

### Lung lavage

This was performed by instilling and collecting 1 ml of PBS through a tracheostomy (20-gauge cannula). Total leukocyte count was determined using a hemocytometer, and differential count by cytocentrifugation of 200 μl of lavage fluid at 300 g for 5 min. Cytospins were stained by Wright-Giemsa for microscopic morphologic cell identification and counting.

### Mucociliary clearance

Mucociliary clearance was measured as the elimination of fluorescent microspheres from the lungs over time. Mice were anesthetized with urethane (2 mg/g, i.p.) and tracheostomized with a blunt beveled 18 gauge Luer stub adapter (Becton, Dickinson). Using a microsprayer (Penn-Century), 25 μl of PBS/0.1% Tween containing 7.43×10^4^ of 4.19 μm fluorescent microspheres (Bangs Laboratories) were loaded at the lung carina through the tracheostomy. Lungs were harvested either immediately (time 0) or mice were mechanically ventilated with a flexiVent (Scireq, Canada) to guarantee uniform ventilation. Lungs were harvested after 30 min, homogenized with 1.5 g of 1.3 mm chrome steel beads (BioSpec) and 1 ml of PBS/0.1% Tween using a Mini-Bead Beater (BioSpec) at 4800 rpm for 3 min. Fluorescent microspheres were then manually counted using a hemocytometer.

### Total bacterial 16S rDNA qPCR

Lungs were harvested and bacterial genomic DNA was extracted and analyzed at Baylor College of Medicine by methods developed for the NIH-Human Microbiome Project (60, 61). Briefly, bacterial genomic DNA was extracted using a PowerSoil DNA Isolation Kit (MO BIO Laboratories, California) following the manufacturer’s instructions. Extracted DNA concentrations were measured by Qubit (Life Technologies) for subsequent normalization of quantitative PCR results (qPCR). qPCR sample analysis was performed in a 7500 Fast Real-Time PCR System. The qPCR primers (1369F-1492R) target regions flanking V9 of the 16S rRNA gene (62). A standard curve was made using a serially diluted plasmid that contains nucleotides 1369 to 1492 of an *E. coli* 16S rRNA gene, and concentrations of the samples were calculated from CT values using the equation generated from plotting the standard curve. All samples were run in triplicate, including the standard curve, a set of non-template controls (NTC), and inhibitor controls (known positives + unknown DNA).

### 16S rRNA gene compositional analysis

The 16S rDNA V4 region was amplified by PCR and sequenced in the MiSeq platform (Illumina) using the 2×250 bp paired-end protocol yielding pair-end reads that overlap almost completely. The primers used for amplification contain adapters for MiSeq sequencing and dual-index barcodes so that the PCR products may be pooled and sequenced directly (63), targeting at least 10,000 reads per sample. The read pairs are demultiplexed based on the unique molecular barcodes, and reads are merged using USEARCH v7.0.1001 (64) allowing zero mismatches and a minimum overlap of 50 bases. Merged reads are trimmed at first base with Q5. In addition, a quality filter is applied to the resulting merged reads, and reads containing above 0.05 expected errors are discarded. 16S rRNA gene sequences were assigned into Operational Taxonomic Units (OTUs) or phylotypes at a similarity cutoff value of 97% using the UPARSE algorithm. OTUs were then mapped to an optimized version of the SILVA Database (65, 66) containing only the 16S v4 region to determine taxonomies. Abundances were recovered by mapping the demultiplexed reads to the UPARSE OTUs.

### Lumenal occlusion in an allergic asthma model

Airway mucus plugging was measured by modifications, as follows, of a method we have described previously (12). Lungs were fixed by immersion, to avoid displacement of lumenal mucus by inflation, in methanol-based Carnoy’s solution (methacarn), to minimize changes in mucus volume, for 48 h at 4°C. Lungs were then excised, and the right caudal lobe (Figure 6, B and figure S6, B), or every lobe (Figure S6, A), was embedded in paraffin. For Figure S6, A and Figure S6, B, a 5 μm section was obtained every 500 μm starting from the most caudal point of the lobe. For Figure 6, B, a 5 μm section was obtained every 1000 μm section of the right caudal lobe, yielding 4 sections per lung. Slides were then deparaffinized, rehydrated, and stained with PAFS. Images were acquired using an upright microscope (Olympus BX 60) with a 20× NA 0.5 lens objective. For quantification, the cross-sectional area of the lumenal mucus was traced (Figure 6, B), and the airway cross-sectional area was also traced to calculate the occlusion fraction (Figure S6, A and figure S6, B), using ImageJ (NIH).

### Lung mechanics

Respiratory resistance was analyzed using a flexiVent system (Scireq). Mice were anaesthetized with urethane (3 mg/g by i.p. injection, a dose sufficient for 2 h of sedation even though experiments last less than 30 min), and paralyzed with succinylcholine chloride (5 mg by i.p. injection followed by continuous i.p. infusion at 20 μg/g·min). Mice were tracheostomized with a blunt beveled 18-gauge Luer-Stub adapter and ventilated at 150 breaths/min, 10 μl/g, against 2-3 cm H_2_O positive end-expiratory pressure. Respiratory resistance was assessed at baseline and in response to four incremental doses of methacholine (MCh) (1, 3, 10 and 30 mg/ml) administered by an in-line ultrasonic nebulizer (4-6 μm, Aerogen, Ireland). Total respiratory resistance was calculated by averaging eight values measured after each dose of MCh for each mouse.

### Lumenal occlusion and emphysema in the β-ENaC-Tg model of cystic fibrosis

Lungs were fixed with 10% NBF and the left lobes were cut in transverse sections starting at the level of the hilum and then every 2 mm from rostral to caudal, yielding 2-4 slices. All slices were embedded in paraffin, and a 5 μm section was obtained from each slice. Sections were then stained with Alcian Blue-Periodic Acid Schiff (AB-PAS) (67). Whole lung section images were obtained in an Olympus BX61VS scanner with a BX81 stage and UPlanSApo 20× NA 0.75 objective lens; scanning conditions were kept constant among specimens. Lumenal occlusion was quantified as in Fig. 5, B from the first 2 sections of each mouse. Emphysema was measured using D_2_, the equivalent mean diameter, computed from measurement of airspace area, and weighted for variance and for skewness towards large spaces (68), from 10 high magnification non-continuous images (1134 × 1134 pixels), equidistant from the center of the section and excluding airways, selected from the sections used in Figure 6, B.

### Statistics

All statistical analyses were performed using GraphPad Prism 7.0 with P<0.05 considered statistically significant. Exact P values and n values for each sample are included in each figure legend. Statistical analysis was performed using one-way ANOVA followed by Tukey’s post hoc test for multiple pair-wise comparisons, and Student’s *t* test or Mann-Whitney *U* test after determining normality of the data using D’Agostino-Pearson omnibus K2 test. A Kolmogorov-Smirnov test was used for Figure S5, D. Values that did not reach significance were not noted. After determining that the expression and secretory function of Munc18 proteins in floxed mice was equivalent to WT, the primary endpoint for studies of secretion, clearance and obstruction was of differences between airway deletant mice and their floxed littermates.

### Study approval

All mice were kept in pathogen-free facilities and handled in accordance with the Institutional Animal Care and Use Committees of The University of Texas MD Anderson Cancer Center, The Texas A&M University Health Science Center Institute of Bioscience and Technology and The University of North Carolina.

## Author contributions

M.J. Tuvim and B.F. Dickey conceived the study. A.M. Jaramillo performed most experiments and analyses. L. Piccotti, W. Velasco, A.S. Huerta, Z. Azzegagh, U. Nazeer, J. Farooq, J. Brenner and J. Parker-Thornburg conducted the remaining experiments. M.J. Tuvim and B.L. Scott generated the conditional Munc18b mouse. R. Adachi generated the conditional Munc18c mouse and helped with statistical and stereological analyses. CM. Evans generated the Muc5ac knockout mouse and assisted with the analysis of mucus occlusion. S. M. Kreda and F. Chung performed the mouse crossing/breeding, the experimental procedures, and data analysis in the β-ENaC-Tg studies. A.R. Burns performed the electron microscopy. A.M. Jaramillo and B.F. Dickey wrote the manuscript. All authors contributed to discussion of the study.

## Acknowledgments

We thank Mehmet Kesimer (UNC-CH) for advice with mucin immunoblots, David E. Ost for assistance with statistical methods, James P. Carson for facilitating use of the program for emphysema analysis, and the MD Anderson Genetically Engineered Mouse Facility (GEMF). For the β-ENaC-Tg studies, we thank the CF Center Molecular Biology Core (Dr. W O’Neal, UNC-CH) for the genotyping, Ms. C van Heusden (UNC-CH) for the BAL preparations, Ms. Kim Burns (UNC-CH) for the lung histology processing and Dr. Hong Dang for the statistical power analysis. A.M.J. thanks the members of her doctoral thesis committee, Magnus Hook, David Reiner, James McNew and Margie Moczygemba, for helpful advice. This work was supported by National Institutes of Health grants R01HL129795, R41HL136057, NIDDK DK065988, R01HL080396, R01HL130938 and R21ES023384; by Cystic Fibrosis Foundation grants DICKEY15P0, DICKEY18G0, KREDA13G0, KREDA16XX0, and BOUCHER15R0 and by Department of Defense grant PR160247. The GEMF at MD Anderson is supported by the Cancer Center Support Grant NCI-CA016672(GEMF). B. L. Scott was supported by the MD Anderson Odyssey Fellowship Program.

